# Astrocytes release prostaglandin E2 to modify respiratory network activity

**DOI:** 10.1101/150250

**Authors:** David Forsberg, Thomas Ringstedt, Eric Herlenius

**Affiliations:** Department of Women’s and Children’s Health, Karolinska Institutet, and Karolinska University Hospital, Stockholm, Sweden

## Abstract

Prostaglandin E2 (PGE2) released during hypercapnic challenge increases Ca^2+^ oscillation frequency in the chemosensitive parafacial respiratory group (pFRG/RTN). Here, we demonstrate that pFRG/RTN astrocytes are the PGE2 source. Two distinct astrocyte subtypes were found using transgenic mice expressing GFP and MrgA1 receptors in astrocytes. Although most astrocytes appeared dormant during time-lapse calcium imaging, a subgroup displayed persistent, rhythmic oscillating calcium activity. These active astrocytes formed a subnetwork within the respiratory network distinct from the neuronal network. Activation of exogenous MrgA1 receptors expressed in astrocytes tripled their calcium oscillation frequency activity in both the preBötzinger complex and pFRG/RTN. However, neurons in the preBötC were unaffected, whereas neuronal calcium oscillatory frequency in pFRG/RTN doubled. Notably, astrocyte activation in pFRG/RTN triggered local PGE2 release and blunted the hypercapnic response. Thus, astrocytes play an active role in respiratory rhythm modulation, modifying respiratory-related behaviorthrough PGE2 release in the pFRG/RTN.

## Introduction

In the last decade, it has become evident that astrocytes are involved in respiratory behavior; electrically active astrocytes are present in the brainstem respiratory center the preBötzinger Complex (preBötC) (Grass, Pawlowski et al. 2004) and exhibit rhythmic calcium (Ca^2+^) oscillations associated with the respiratory-related neuronal rhythm (Schnell, Fresemann et al. 2011, Okada, Sasaki et al. 2012). Further, brainstem astrocytes respond to changes in blood gas levels (Angelova, Kasymov et al. 2015), especially in the chemosensitive region the parafacial respiratory group/retrotrapezoid nucleus (pFRG/RTN) (Huckstepp, Bihi et al. 2010). During the hypercapnic response, carbon dioxide (CO_2_) causes a vesicle-dependent release of ATP from astrocytes (Fukuda, Honda et al. 1978, Erlichman, Leiter et al. 2010, Gourine, Kasymov et al. 2010, Huckstepp, Bihi et al. 2010, Turovsky, Karagiannis et al. 2015, Turovsky, Theparambil et al. 2016). ATP may also be released in a vesicle independent manner through CO_2_-sensitive gap junctions (Huckstepp, Bihi et al. 2010, Huckstepp, Eason et al. 2010, Meigh, Greenhalgh et al. 2013). ATP then increases inspiratory frequency through actions on P2 receptors located in neurons in both the pFRG/RTN and the preBötC (Lorier, Huxtable et al. 2007, Gourine, Kasymov et al. 2010). Acting via EP3 receptors, prostaglandin E2 (PGE2) also alters respiratory network activity by inhibiting signaling frequency in the preBötC while increasing it in the pFRG/RTN (Forsberg, Horn et al. 2016). Additionally, the response to hypercapnic and hypoxic challenges are blunted by PGE2 (Hofstetter, Saha et al. 2007, Siljehav, Olsson Hofstetter et al. 2012, Siljehav, Hofstetter et al. 2015). Because PGE2 is released through gap junctions during hypercapnia, and since some astrocytes in the pFRG/RTN express the primary enzyme involved in PGE2 synthesis (i.e., mPGEs-1) (Forsberg, Horn et al. 2016), we hypothesized that hypercapnia induces astrocytes to release PGE2 in addition to ATP. Thus, astrocytes may also modulate respiratory-related network rhythms via the downstream actions of PGE2.

To investigate the role of astrocytes in respiratory rhythm-generating networks and determine whether astrocytes release PGE2, we utilized organotypic brainstem slice cultures (Forsberg, Horn et al. 2016, Phillips, Herly et al. 2016) of B6.Cg-Tg(hGFAP-tTA:: tetO-MrgA1) 1^Kdmc/Mmmh^ mice (GFAP^MrgA1+^) in which green fluorescent protein (GFP) and the MrgA1 receptor (MrgA1R) were expressed under control of the promoter for glial fibrillary acidic protein (GFAP), a generally used marker for astrocytes (Brenner, Kisseberth et al. 1994). The MrgA1R is an endogenous G_q_-coupled receptor that is normally not expressed in the brain. Rather, MrgA1R is expressed exclusively in the dorsal root ganglion nociceptive sensory terminals of the spinal cord (Fiacco, Agulhon et al. 2007) and is activated by RF peptides (Dong, Han et al. 2001). Consequently, we could selectively activate brainstem astrocytes using synthetic Phe-Leu-Arg-Phe-amide (FLRF) (Young, Platel et al. 2010, Cao, Li et al. 2013).

## Results and Discussion

Expression of GFP in the pFRG/RTN and preBötC of the GFAP^MrgA1+^ mice was evident (**Figure 1**), and GFP was co-localized with GFAP immunolabeling (**Figure 1b and h**, n = 11 slices; see **Table 1** for the number of times each experiment was conducted). Although subgroups of GFAP-expressing cells have been previously defined using fluorescence intensity (Grass, Pawlowski et al. 2004), we were unable to detect measurable differences in the GFP fluorescence among our samples. The astrocyte and oligodendrocyte marker S100β was expressed by 12% ± 3% of the GFP-positive cells (n = 13 slices). Conversely, 92% ± 4% of the S100β-positive cells also expressed GFP (**Figure 1c and i,** n = 13 slices). The GFP-positive cells followed the neuronal spread but expressed neither the neuronal markers NK1R or MAP2 (**Figure 1d, e and j, k,** n = 21 slices) nor the microglial marker Iba1 (**Figure 1f and l**, n = 11 slices). No GFP expression was detected in littermate controls (WT; **Figure 1m**, n = 12 slices). These findings confirm that the GFAP-driven expression of GFP (and thus MrgA1R) is astrocyte specific, as expected (Brenner, Kisseberth et al. 1994, Fiacco, Agulhon et al. 2007, Cao, Li et al. 2013). However, the antibodies in our immunohistochemistry experiments penetrated only approximately 20% of the brainstem slice thickness (**Figure 1g**), which must be considered when the organotypic slice culture method is used.

**Figure 1.**
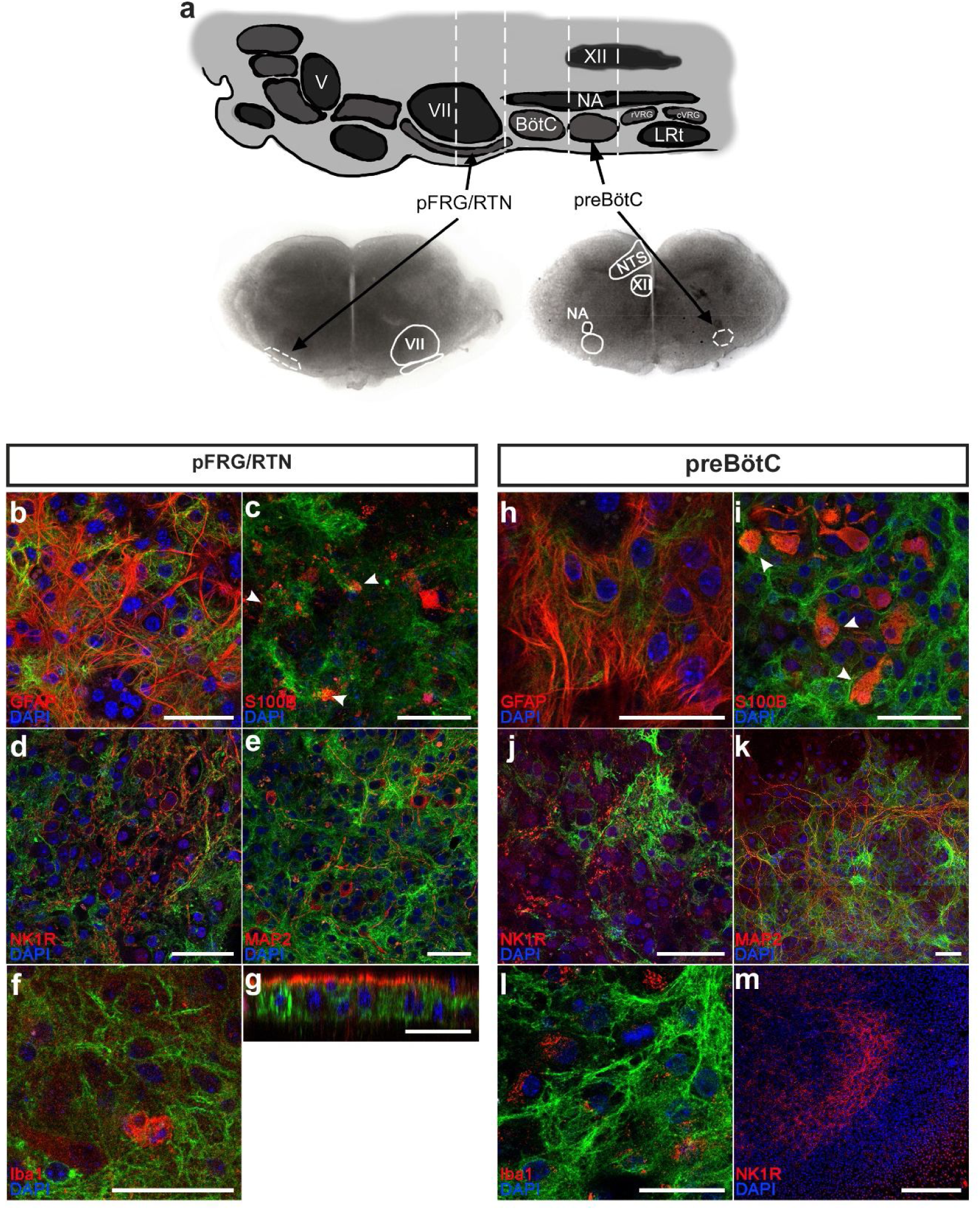
Astrocytes identified with GFP. The pFRG/RTN and preBötC slices (a) obtained from GFAP^MrgA1+^ mice express GFP (green), which co-localizes with the astrocytic markers GFAP (red; b, h) and S100β (c, i). By contrast, no GFP co-expression is detected with the neuronal markers NK1R (red; d, j) or MAP2 (red; e, k), or with the microglial marker Iba1 (red; f, l). Littermate controls do not express GFP (m). The z-projections revealed that the antibodies used in the immunohistochemical analyses penetrate only the outer 20% of the brainstem slice (g). V; trigeminal nucleus, VII; facial nucleus; BötC; Bötzinger complex, NA; Nucleus Ambiguus, XII; hypoglossal nucleus, rVRG; rostral ventral respiratory group, cVRG; caudal ventral respiratory group, LRt; lateral reticular nucleus, NTS; nucleus tractus solitaries. Arrowheads indicate double-labeled cells. Scale bars: 50 μm in b–l, 500 μm in m.

**Table 1.** Number of experiments conducted.

| *Figure* | *Panel* | *Number of experiments* |
| --- | --- | --- |
| 1 | b | 5 |
|  | c | 7 |
|  | d | 6 |
|  | e | 6 |
|  | f | 6 |
|  | g | 5 |
|  | h | 6 |
|  | i | 6 |
|  | j | 5 |
|  | k | 5 |
|  | l | 5 |
|  | m | 6 |
| 3 | a | 19 |
|  | b | 22 |

The Ca^2+^ time-lapse imaging of brainstem slices derived from both GFAP^MrgA1+^ and WT control mice displayed similar average Ca^2+^ signaling frequency and network structure. Thus, the induced expression of GFP and MrgA1R did not affect respiratory network activity or function. Taken together with the results of the immunohistochemical expression analyses, this demonstrates that the MrgA1 mouse line has astrocyte-specific expression of the inserted genes, which display no physiological effects, making this mouse line useful for investigating the role of astrocytes in respiratory networks.

We examined the Ca^2+^-signaling activity of cells in the pFRG/RTN and in the preBötC and identified two subgroups among the GFP-expressing astrocytes: cells with a rhythmic Ca^2+^-signaling pattern, and cells that appeared inactive, that is, the Fura-2 fluorescence intensity in these cells was stable over time. A similar subgrouping of astrocytes, with approximately 10% exhibiting calcium transients preceding inspiratory-related neuronal signals, has been suggested previously on the basis of both electrophysiological and Ca^2+^ imaging methods (Grass, Pawlowski et al. 2004, Schnell, Fresemann et al. 2011, Oku, Fresemann et al. 2016). The present study also found that the majority of the astrocytes appeared inactive (**Figure 2a and b**; 82% ± 9% in the pFRG/RTN, n = 19 slices; 87% ± 7% in the preBötC, n = 22 slices). In the preBötC, the proportion of active astrocytes was similar to that detected by Schnell and colleagues (Schnell, Fresemann et al. 2011), but lower than that reported by both Grass and colleagues (Grass, Pawlowski et al. 2004) and Oku and colleagues (Oku, Fresemann et al. 2016). However, our data are presented as the proportion per respiratory network, whereas previous studies evaluated the proportion of the total number of recorded cells, independent of individual networks. To our knowledge, the present study is the first to describe active and inactive astrocytes within the pFRG/RTN. Notably, we observed the following difference between the pFRG/RTN and the preBötC: 40% ± 12% of the active cells in pFRG/RTN were astrocytes, whereas only 20% ± 9% of active cells in the preBötC were astrocytes (**Figure 2a and b**, p < 0.05 when comparing the pFRG/RTN and the preBötC, n = 41 slices; see **Table 2** for a list of all statistical tests used for all comparisons in the present study).

**Figure 2.**
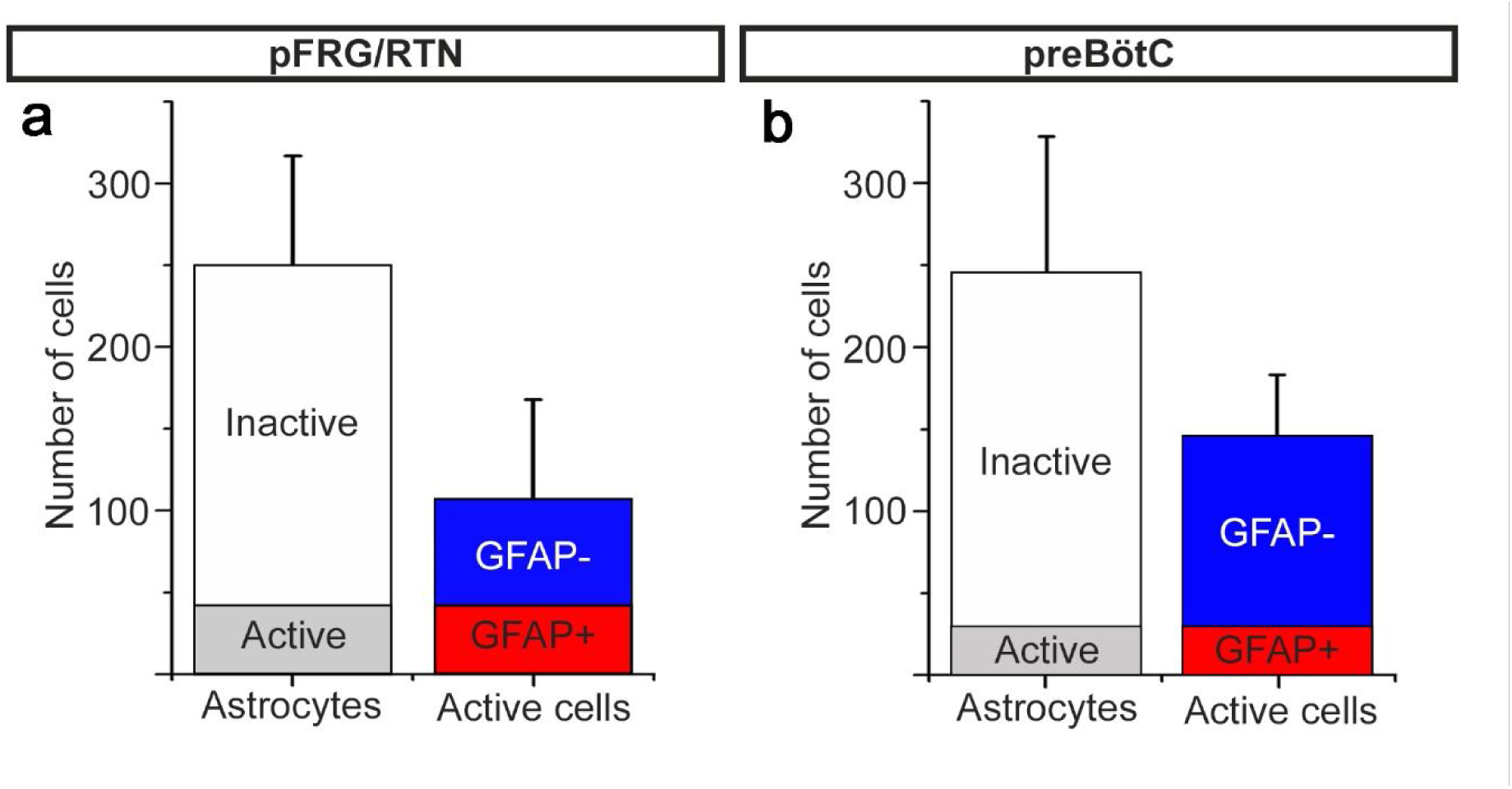
Active astrocytes constitute a subgroup of glial cells within respiratory networks. In the pFRG/RTN, 18% ± 13% of the astrocytes are active. Of the total number of active cells, 40% ± 12% are astrocytes (GFAP+) (a). In the preBötC, 13% ± 7% of the astrocytes are active, and only 20% ± 9% of the total number of active cells are astrocytes (b). Data are presented as the mean ± SD.

**Table 2.** Statistical analyses and results.

| *Figure* | *Test used* | *Exact p-value* | *Degrees of freedom*<br>*&*<br>*F/t/z/R value* |
| --- | --- | --- | --- |
| 2 | Student's t-test | 0.0261 | DF = 40 |
| 3c | Full factorial ANOVA | 0.821<br>0.688<br>0.716 | F=0.071<br>F=0.043<br>F=0.052<br>DFE=18 |
| 3e | Full factorial ANOVA | 0.416<br>0.237<br>0.0946 | F=0.064<br>F=0.065<br>F=0.038<br>DFE=21 |
| 4b | Student's t-test | 0.0213<br>0.0278<br>0.0342<br>0.0413<br>0.0427 | DF=10, 10, 10, 7, 7 |
|  |  | 0.0147<br>0.0129<br>0.0385<br>0.0311<br>0.0406 | DF=11, 11, 11, 9, 9 |
| 4c | Full factorial ANOVA | 0.416<br>0.759 | F=0.029, DFE=6<br>F=0.063, DFE=7 |
| 5a | Student's t-test | 0.0729<br>0.0271<br>0.0014<br>0.0298<br>0.194 | DF=7 |
| 5b | Student's t-test | 0.017, 0.44<br>0.014, 0.71<br>0.29, 0.024 | DF=8 |
| 5c | Student's t-test | 0.018, 0.41<br>0.74, 0.85<br>0.66, 0.70 | DF=7 |

On the basis of a correlated cell activity analysis (Smedler, Malmersjo et al. 2014, Forsberg, Horn et al. 2016), we determined that the active astrocytes within the pFRG/RTN and within the preBötC formed specific networks. These networks differed from those of non-astrocytes, that is, neurons (**Figure 3a and b;** see **Table 1** for the number of times each experiment was conducted), but displayed similar network structures (**Figure 3c and e**). The astrocytic and neuronal networks connected with each other at multiple nodes, and together they formed the respiratory networks of the pFRG/RTN and of the preBötC. In the pFRG/RTN, astrocytes constituted 37% ± 13% of the total network (number of correlating cell pairs), whereas astrocytes made up only 16% ± 5% of the preBötC network (p < 0.05 when comparing the pFRG/RTN and the preBötC; **Figure 3a, d, b and f**). We also observed a significantly larger number of connections between astrocytes and neurons in the pFRG/RTN than in the preBötC (33% ± 12% vs. 24% ± 11 % of all correlating cell pairs, p < 0.05; **Figure 3a, b, d and f**). Thus, astrocytes are both independently active and part of the respiratory networks, but appear to have different actions in the two central pattern generators.

**Figure 3.**
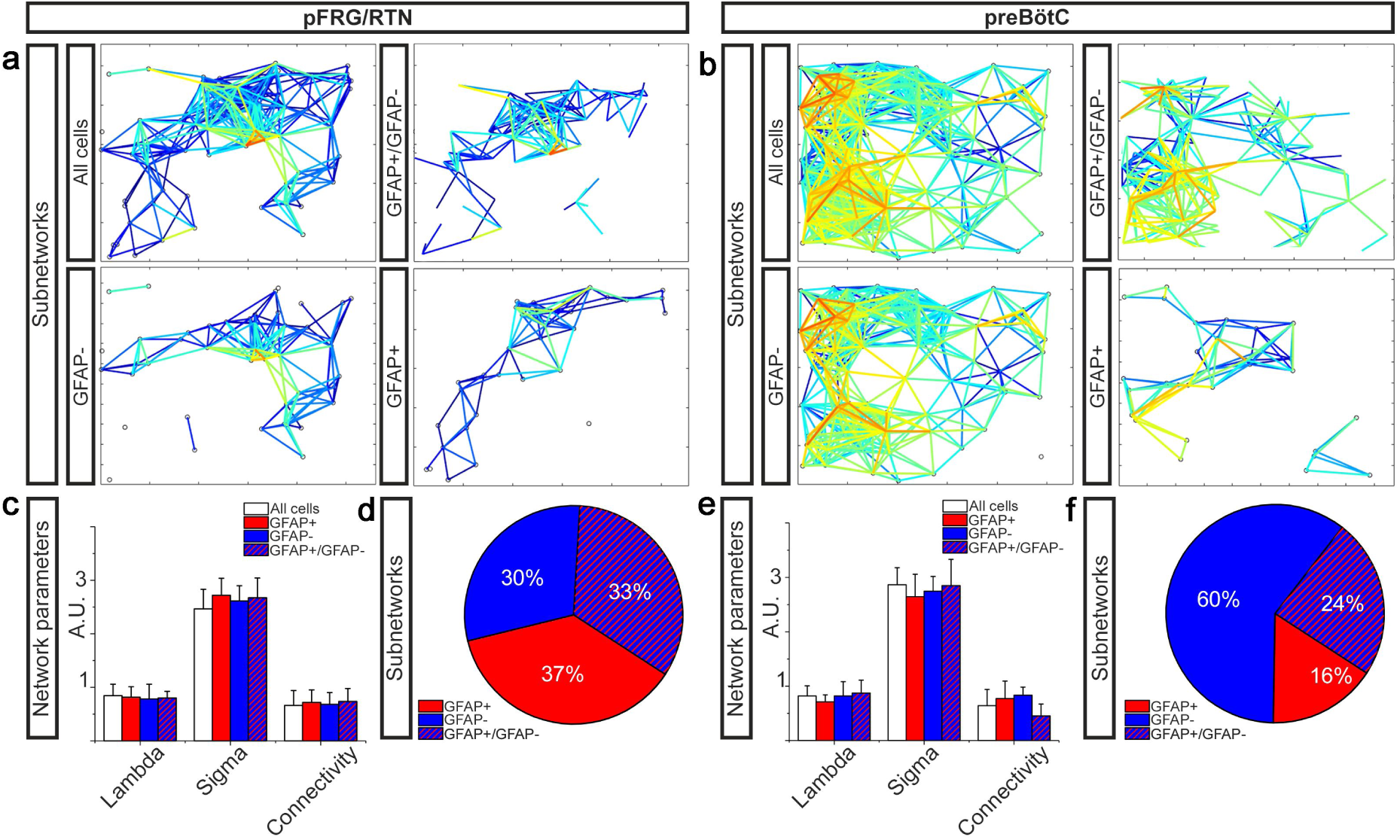
Astrocytic and neuronal networks are separate but collaborate to build respiratory networks. (a) and (b) graphically represent the network structures in a 2-D plane, with the lines representing correlation coefficients above set cut-offs for the cell pairs (warmer color equals higher correlation coefficient). In both the pFRG/RTN and the preBötC, active astrocytes (GFAP+) and neurons (GFAP-) form separate networks (a, b). However, these networks interconnect to build a joint astrocyte–neuron (GFAP+/GFAP-) network (a, b). Together, this interconnected network is the whole respiratory network (a, b). All subnetworks have similar network properties (c, e), except the connectivity of the astrocyte–neuron network in the preBötC, which is slightly and nonsignificantly less than that of the other subnetworks (e). The pFRG/RTN consists of equal parts of the three subnetworks (d), whereas the preBötC network is predominantly neuronal (f). Data are presented as the mean ± SD.

Under control conditions, astrocytes displayed a low oscillating frequency (19 ± 8 mHz in the preBötC (n = 22 slices) vs. 18 ± 7 mHz in the pFRG (n = 19 slices); difference not statistically significant), similar to that described previously for active astrocytes (Grass, Pawlowski et al. 2004, Schnell, Fresemann et al. 2011, Okada, Sasaki et al. 2012, Oku, Fresemann et al. 2016). The frequency displayed by the astrocytes in each region was lower than that shown by the neurons in the respective regions (preBötC: 90 ± 40 mHz, n = 22 slices, p < 0.05 compared with astrocytes; pFRG: 106 ± 58 mHz, n = 19 slices, p < 0.05 compared with astrocytes). We did not register nerve output signals in the present study and can therefore not evaluate whether astrocytic signaling was phase-locked with the respiratory output.

Next, we activated the GFAP^MrgA1+^ astrocytes by applying the MrgA1R ligand FLRF, which increased the Ca^2+^-signaling frequency of astrocytes in both the pFRG/RTN and the preBötC (**Figure 4a and b**). In the pFRG/RTN, astrocyte activation also induced an increase in the signaling frequency of the non-astrocytes (**Figure 4a and b**). This is in contrast to the preBötC, wherein non-astrocytes retained their Ca^2+^-signaling frequency independent of astrocyte stimulation (**Figure 4a and b**). Inactive astrocytes showed a single Ca^2+^ peak shortly after the application of FLRF, but returned to the inactive state within 10 s (**Figure 4a**). The stimulation of astrocytes did not affect the network structures of the astrocytic, neuronal, or complete respiratory networks (**Figure 4c**). Astrocytes in slices derived from WT mice did not react to the application of FLRF (**Figure 4b**).

**Figure 4.**
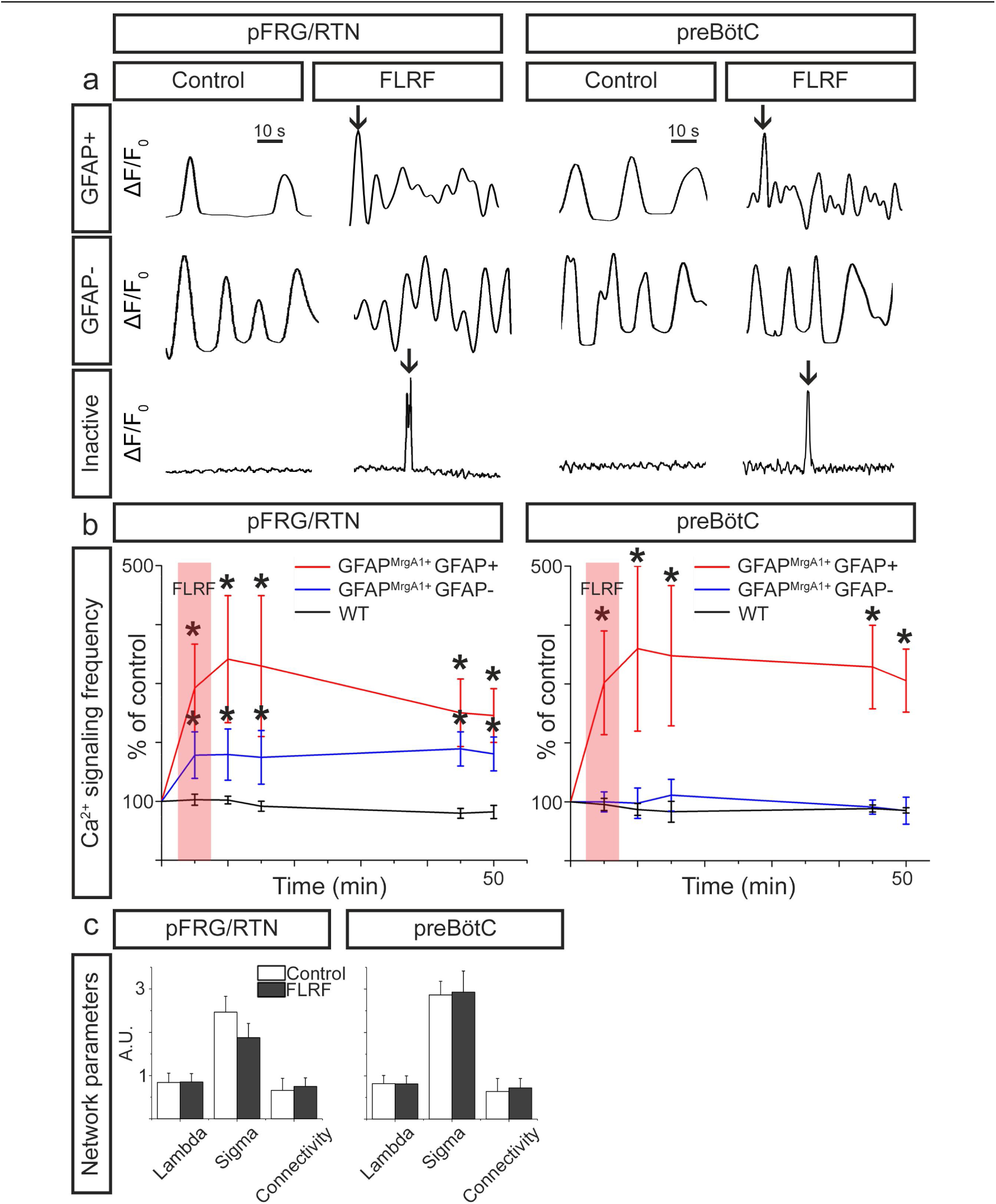
Astrocytes control neurons in the pFRG/RTN. (a) Individual 1 min calcium traces (ΔF/F_0_ over time, band pass filtered 0.01–0.15 Hz) during the control period and after the application of the MrgA1R ligand FLRF. Activation of astrocytes through addition of FLRF increases calcium-signaling frequency of astrocytes in both the pFRG/RTN and the preBötC (b, red trace). In the pFRG/RTN, neuronal calcium-signaling frequency increases after astrocyte activation (b, blue trace). This is not observed in the preBötC (b, blue trace). Littermate controls (WT) do not respond to FLRF (b, black trace). Network properties are not affected by addition of FLRF (c). Data are presented as the mean ± SD. *p < 0.05 compared to their respective controls.

While these findings suggest that astrocytes do not directly modify neuronal activity in the preBötC, astrocytes may, nonetheless, be important for the maintenance of the respiratory rhythm generation in this complex. For instance, the astrocyte inhibitor methionine sulfoximine depresses breathing in vivo (Young, Dreshaj et al. 2005), and glial inhibitors (fluorocitrate, fluoroacetate, and methionine sulfoximine) reduce respiratory-related activity in the preBötC in vitro (Erlichman, Li et al. 1998, Huxtable, Zwicker et al. 2010). In addition, hypoxia can induce a Ca^2+^ influx in preBötC astrocytes, inducing a release of ATP (Gourine, Llaudet et al. 2005, Angelova, Kasymov et al. 2015). This helps to maintain the inspiratory-related rhythm generation in the preBötC during post-hypoxic depression (Gourine, Llaudet et al. 2005). Although we did not measure an increase in neuronal network burst frequency after astrocyte-specific activation, we did not test responses under hypoxic conditions.

Our data suggest that, in contrast to their function in the preBötC, astrocytes in the pFRG/RTN modulate respiratory network activity. This corroborates the results of previous studies that have demonstrated the involvement of astrocytes in chemosensitivity through purinergic signaling (Erlichman, Leiter et al. 2010, Gourine, Kasymov et al. 2010, Huckstepp, Bihi et al. 2010, Gourine and Kasparov 2011, Angelova, Kasymov et al. 2015). In one of our recent articles, we suggested that astrocytes release PGE2 (in addition to ATP) in response to a hypercapnic challenge (Forsberg, Horn et al. 2016). This would in part explain the interaction between chemosensitivity and the inflammatory system observed in several studies (Hofstetter, Saha et al. 2007, Siljehav, Olsson Hofstetter et al. 2012, Siljehav, Shvarev et al. 2014, Forsberg, Horn et al. 2016). Moreover, in human neonates, rapid elevation of brainstem PGE2 during infectious events is associated with and may explain the initial presenting symptoms of infection, which are apnea, bradycardia, and desaturation (Siljehav, Hofstetter et al. 2015).

Because a hypercapnic challenge triggers a gap junction mediated release of PGE2 (Forsberg, Horn et al. 2016), we hypothesized that astrocyte stimulation would trigger a similar release. Indeed, we found that after FLRF application, the PGE2 levels in the artificial cerebrospinal fluid (aCSF) doubled in the pFRG/RTN (**Figure 5a**, n = 8 slices, p < 0.05). The increase was transient, similar to the PGE2 release that occurs during hypercapnia (Forsberg, Horn et al. 2016). By contrast, the PGE2 levels remained unaffected by FLRF application in the GFAP^MrgA1+^ preBötC slices (n = 3 slices) and both the pFRG/RTN and preBötC regions in the WT slices (n = 2 slices for each region). These results indicated that the pFRG/RTN contains chemosensitive astrocytes that release PGE2 upon hypercapnic challenge. The physiological effect of the released PGE2 is likely multifactorial. We previously demonstrated that PGE2 is involved in the stimulation of the pFRG/RTN (Forsberg, Horn et al. 2016), similar to the effects of ATP (Gourine, Kasymov et al. 2010). Although ATP counteracts the vasodilatory effect of CO_2_/H^+^ during hypercapnia in the ventral brainstem (Hawkins, Takakura et al. 2017), the role of PGE2 in the microcircuits controlling brainstem blood flow will require further investigation.

**Figure 5.**
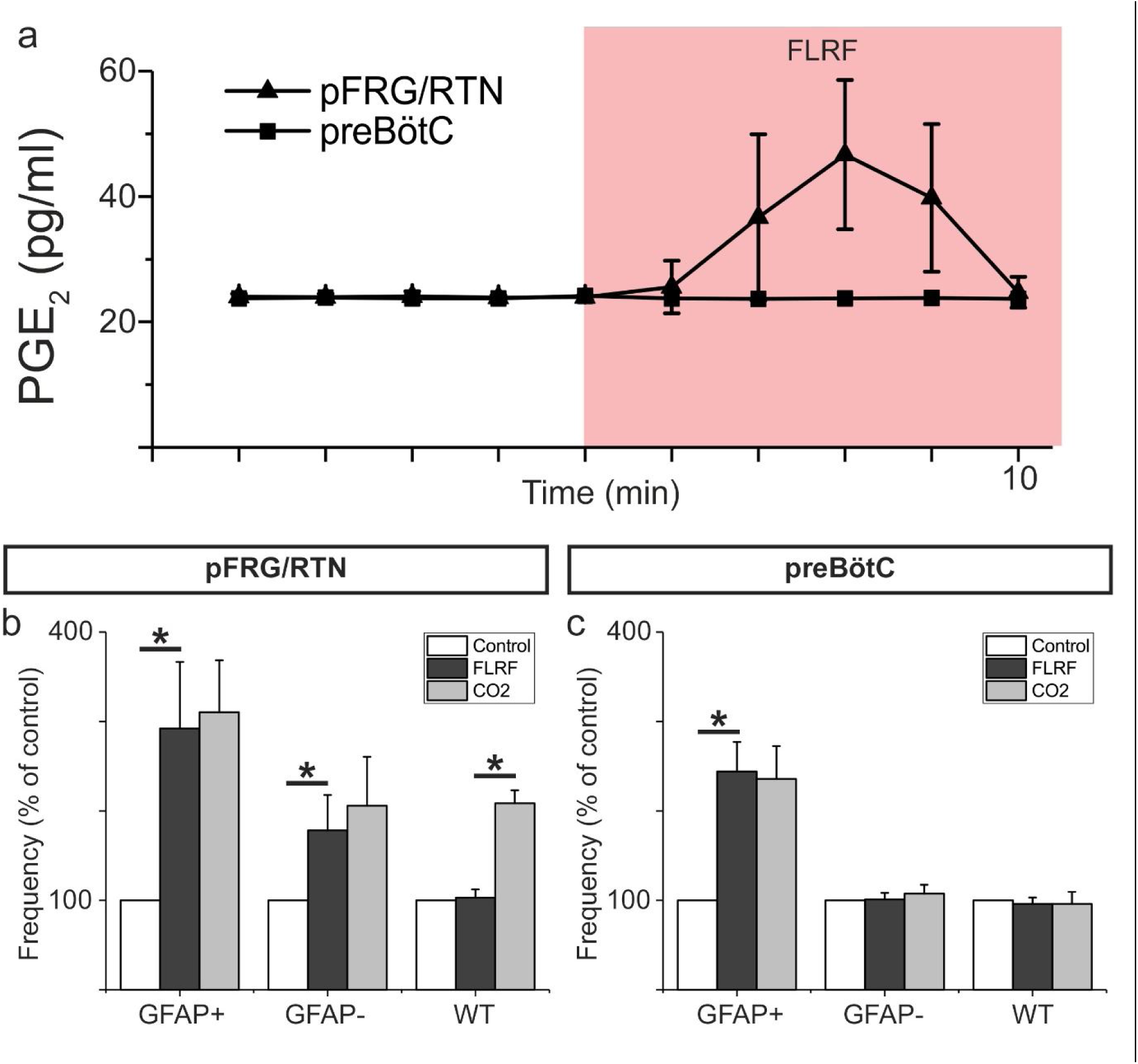
Astrocytes release PGE2 upon activation. Astrocyte activation through application of the MrgA1R ligand FLRF increases PGE2 levels in aCSF (a). After such activation, the hypercapnic response (CO_2_ partial pressure increase from 4.6 kPa to 6.6 kPa) is blunted (b). The preBötC does not respond to a hypercapnic challenge (c). Data are presented as the mean ± SD. *p < 0.05 for the indicated comparisons.

We also determined that pre-activation of astrocytes blunts the hypercapnic response in the pFRG/RTN (**Figure 5b**, n = 8 slices). This result suggested that existing PGE2, and likely other gliotransmitters, is released after activation, depleting the stores. Extracellular pH is retained during hypercapnia in our system (Forsberg, Horn et al. 2016), but CO_2_ can pass over the cell membrane and decrease the intracellular pH. This decreased pH drives bicarbonate and sodium ions into the cell, triggering a Ca^2+^ influx (Turovsky, Theparambil et al. 2016). In parallel, CO_2_ can directly modify connexin 26 hemichannels to induce an open state (Huckstepp, Bihi et al. 2010, Meigh, Greenhalgh et al. 2013). Thus, it is possible that the mechanism behind the gliotransmitter release during hypercapnic challenge is different from that of the FLRF-induced activation and Ca^2+^ influx. The design of the present study could not discern between PGE2 released from active and inactive astrocytes or determine whether PGE2 directly affected neurons or acted via intermediate astrocytes. In the pFRG/RTN, prostaglandin EP3 receptors have been found on both neurons and astrocytes (Forsberg, Horn et al. 2016). Thus, the different pathways as well as the kinetics associated with the effects of PGE2 will require further investigation. Hypercapnia did not affect the preBötC activity (**Figure 5c**, n = 6 slices), consistent with the results of our previous study (Forsberg, Horn et al. 2016). In summary, these results indicated that the pFRG/RTN contains astrocytes that are able to react to CO_2_ and release PGE2 (and possibly other gliotransmitters) to modify the behavior of the neuronal population. This type of astrocytic responsiveness and modifying effect was not detected in the preBötC.

Taken together, our results led us to conclude that a subgroup of astrocytes with oscillatory Ca^2+^ activity interacted with and was an essential part of the respiratory neural networks. We observed that almost half of the cells in the pFRG/RTN network were astrocytes and that these astrocytes could entrain the neuronal rhythm, suggesting that pFRG/RTN astrocytes are a source of gliotransmitters that modify respiratory activity. Moreover, a transient release of PGE2 induced by a Ca^2+^ influx in the astrocytes of the pFRG/RTN substantially reduced the subsequent hypercapnic response, suggesting a central role of PGE2 as a gliotransmitter in chemosensitivity. Thus, subgroups of astrocytes participate in the respiratory rhythm of both the preBötC and the pFRG/RTN, and modify respiratory network behavior in the pFRG/RTN. Therefore, we suggest that astrocytes constitute an important link between the respiratory and inflammatory systems, and stand out as a potential target for the treatment of central respiratory dysfunction.

## Materials and Methods

### Subjects

Mice expressing green fluorescent protein (GFP) under the GFAP promoter were used. Frozen sperm from the GFAP-tTA (Lin, Kemper et al. 2004, Pascual, Casper et al. 2005) and tetO-Mrgpra1 (Fiacco, Agulhon et al. 2007) mouse strains were purchased from the Mutant Mouse Resource and Research Center (MMRRC) supported by the National Institutes of Health. The strains were re-derived by the Karolinska Center for Transgene Technologies, and the offspring were crossed as previously described (Fiacco, Agulhon et al. 2007). The double transgenic B6.Cg-Tg(hGFAP-tTA::tetO-MrgA1)^1Kdmc/Mmmh^ mice were identified by polymerase chain reaction analyses according to instructions provided by the MMRRC.

All mice were reared by their mothers under standardized conditions with a 12 h light–dark cycle. The mice were allowed food and water ad libitum. The studies were performed in accordance with European Community Guidelines and approved by the regional ethics committee. The animals were reared and kept in the Department of Comparative Medicine at the Karolinska Institute in Stockholm, Sweden.

### Brainstem organotypic culture

Mouse pups were used at postnatal day 3 for the establishment of brainstem organotypic slice cultures (Herlenius, Thonabulsombat et al. 2012, Forsberg, Horn et al. 2016). Transverse slices (300 μm thick) were maintained in culture for 7 to 14 days. Slices were selected by using online anatomical references (Ruangkittisakul, Schwarzacher et al. 2006, Ruangkittisakul, Panaitescu et al. 2011, Ruangkittisakul, Kottick et al. 2014).

### Immunohistochemistry

The immunohistochemistry procedure was the same as that described previously (Forsberg, Horn et al. 2016).). The primary antibodies used were mouse anti-microtubule associated protein 2 (MAP2; Invitrogen, cat. no. P11137), rabbit anti-neurokinin 1 receptor (NK1R; Sigma-Aldrich, St. Louis, MO, USA, cat no. S8305), mouse anti-GFAP (Chemicon, Temecula, CA, USA, cat no. MAB360), rabbit anti-S100β (Millipore; cat. no. 04-1054), rabbit anti-Iba1 (Wako, Japan; cat. no. 019-19741) and goat anti-GFP (Abcam, Cambridge, United Kingdom; cat. no. Ab6673). The secondary antibodies used were Alexa Fluor Plus 555 goat anti-mouse IgG (Thermo Fisher Scientific, Waltham, MA, USA; cat. no. A32737), Alexa Fluor Plus 555 goat anti-rabbit IgG (Thermo Fisher Scientific, Waltham, MA, USA; cat. no. A32732), Alexa Fluor 555 donkey anti-rabbit (Thermo Fisher Scientific, Waltham, MA, USA; cat. no. A31572) and Alexa Fluor 488 donkey anti-goat (Thermo Fisher Scientific, Waltham, MA, USA; cat. no. A11055). Negative control tissue incubated in the absence of primary antibodies showed no staining.

### Ca^2+^ time-lapse imaging

For Ca^2+^ imaging, Fura-2 AM (Thermo Fisher Scientific, Waltham, MA, USA; cat. no. F1201) dissolved in DMSO (Sigma-Aldrich, St. Louis, MO, USA, cat. no .D2650) was used at 166 μM in aCSF (containing in mM: 150.1 Na^+^, 3 K^+^, 2 Ca^2+^, 2 Mg^2+^, 135 Cl^−^, 1.1 H_2_PO_4_^−^, 25 HCO_3_^-^ and 10 glucose) together with 0.02% pluronic acid (Thermo Fisher Scientific, Waltham, MA, USA cat. no. P3000MP). To localize the preBötC or the pFRG/RTN, tetramethylrhodamine-conjugated substance P (TMR-SP; Biomol, Oakdale, NY, USA) was used at a final concentration of 3 μM in aCSF. The TMR-SP solution was placed on top on the brainstem slice and incubated for 10 min at 37°C in an atmosphere of 5% CO_2_. Brainstem slices were incubated in the TMR-SP solution for 10 min at 37°C in an atmosphere of 5% CO_2_. The TMR-SP solution was then replaced with 1.5 ml of 166 μM Fura-2 solution. The Fura-2 solution was incubated for 30 min at room temperature. Before imaging, the slice was washed with aCSF for 10 min (32°C, 5% CO_2_).

During time-lapse imaging, slices were kept in an open chamber perfused with aCSF (1.5 mL/min), as described previously (Forsberg, Horn et al. 2016). The exposure time was set to 100 ms, with an imaging interval of 0.5 s. During imaging, FLRF (10 μM in 0.1% DMSO in aCSF; Innovagen, Lund, Sweden) was added continuously for 5 min after a control period. A subset of slices was exposed to isohydric hypercapnia, with the aCSF adjusted with a high-bicarbonate buffer concentration (in mM: 150.1 Na^+^, 3 K^+^, 2 Ca^2+^, 2 Mg^2+^, 111 Cl^−^, 1.1 H_2_PO_4_^−^, 50 HCO_3_^−^, and 10 glucose) saturated with 8% CO_2_. This generated a hypercapnic carbon dioxide partial pressure (pCO_2_) of 6.6 kPa at pH 7.5.

### PGE_2_ enzyme-linked immunosorbent assay (ELISA)

The release of PGE2 in aCSF during FLRF exposure was assessed by ELISA. The aCSF samples were collected through the perfusion system each minute and stored at −80°C. The prostaglandin E2 EIA monoclonal kit (Cayman Chemicals, Ann Arbor, MI, USA, cat. no. 514010) was used according to our previously published procedure (Forsberg, Horn et al. 2016).

### Data analysis

Data were analyzed as previously described (Forsberg, Horn et al. 2016).

### Statistics

Statistical analysis of paired comparisons was performed by Student’s *t*-test. Full factorial analysis of variance (ANOVA) was performed when more than one independent variable was compared or for multiple comparisons. Both tests were two-sided. The compared data were of equal variance and normally distributed. All calculations for the statistical tests were conducted with JMP software (version 11.1., SAS Institute Inc., Cary, NC, USA). In all cases, values of p <0.05 were considered statistically significant. Data are presented as means ± SD. All data sets were compared less than 20 times; thus no statistical corrections were applied. Because these experiments were conducted to provide new descriptive data, no explicit power analysis was performed. Instead, sample sizes similar to previous publications using similar methods were used.

## Acknowledgments

We thank Torkel Mattesson, Wiktor Phillips and Evangelia Tserga for technical assistance. This study was supported by the Swedish Research Council, the Stockholm County Council, the Karolinska Institutet, and grants from the Brain Foundation, M & M Wallenberg, Fraenkel, Axel Tielman, Freemasons Children’s House and Swedish National Heart and Lung Foundations.

## References

Angelova PR, Kasymov V, Christie I, Sheikhbahaei S, Turovsky E, Marina N, Korsak A, Zwicker J, Teschemacher AG, Ackland GL, Funk GD, Kasparov S, Abramov AY and Gourine AV. 2015. Functional Oxygen Sensitivity of Astrocytes. J Neurosci 35:10460-10473. 10.1523/JNEUROSCI.0045-15.2015.

Brenner M, Kisseberth WC, Su Y, Besnard F and Messing A. 1994. GFAP promoter directs astrocyte-specific expression in transgenic mice. J Neurosci 14:1030-1037.

Cao X, Li LP, Wang Q, Wu Q, Hu HH, Zhang M, Fang YY, Zhang J, Li SJ, Xiong WC, Yan HC, Gao YB, Liu JH, Li XW, Sun LR, Zeng YN, Zhu XH and Gao TM. 2013. Astrocyte-derived ATP modulates depressive-like behaviors. Nat Med 19:773-777. 10.1038/nm.3162.

Dong X, Han S, Zylka MJ, Simon MI and Anderson DJ. 2001. A diverse family of GPCRs expressed in specific subsets of nociceptive sensory neurons. Cell 106:619-632.

Erlichman JS, Leiter JC and Gourine AV. 2010. ATP, glia and central respiratory control. Respir Physiol Neurobiol 173:305-311. S1569-9048(10)00227-2 [pii] 10.1016/j.resp.2010.06.009.

Erlichman JS, Li A and Nattie EE. 1998. Ventilatory effects of glial dysfunction in a rat brain stem chemoreceptor region. J Appl Physiol (1985) 85:1599-1604.

Fiacco TA, Agulhon C, Taves SR, Petravicz J, Casper KB, Dong X, Chen J and McCarthy KD. 2007. Selective stimulation of astrocyte calcium in situ does not affect neuronal excitatory synaptic activity. Neuron 54:611-626. S0896-6273(07)00336-4 [pii] 10.1016/j.neuron.2007.04.032.

Forsberg D, Horn Z, Tserga E, Smedler E, Silberberg G, Shvarev Y, Kaila K, Uhlen P and Herlenius E. 2016. CO_2_-evoked release of PGE2 modulates sighs and inspiration as demonstrated in brainstem organotypic culture. Elife 5:e14170. 10.7554/eLife.14170.

Fukuda Y, Honda Y, Schlafke ME and Loeschcke HH. 1978. Effect of H+ on the membrane potential of silent cells in the ventral and dorsal surface layers of the rat medulla in vitro. Pflugers Arch 376:229-235.

Gourine AV and Kasparov S. 2011. Astrocytes as brain interoceptors. Exp Physiol 96:411-416. 10.1113/expphysiol.2010.053165.

Gourine AV, Kasymov V, Marina N, Tang F, Figueiredo MF, Lane S, Teschemacher AG, Spyer KM, Deisseroth K and Kasparov S. 2010. Astrocytes control breathing through pH-dependent release of ATP. Science 329:571-575. science.1190721 [pii] 10.1126/science.1190721.

Gourine AV, Llaudet E, Dale N and Spyer KM. 2005. Release of ATP in the ventral medulla during hypoxia in rats: role in hypoxic ventilatory response. J Neurosci 25:1211-1218. 10.1523/JNEUROSCI.3763-04.2005.

Grass D, Pawlowski PG, Hirrlinger J, Papadopoulos N, Richter DW, Kirchhoff F and Hulsmann S. 2004. Diversity of Functional Astroglial Properties in the Respiratory Network. J. Neurosci. 24:1358-1365.

Hawkins VE, Takakura AC, Trinh A, Malheiros-Lima MR, Cleary CM, Wenker IC, Dubreuil T, Rodriguez EM, Nelson MT, Moreira TS and Mulkey DK. 2017. Purinergic regulation of vascular tone in the retrotrapezoid nucleus is specialized to support the drive to breathe. Elife 6. 10.7554/eLife.25232.

Herlenius E, Thonabulsombat C, Forsberg D, Jaderstad J, Jaderstad LM, Bjork L and Olivius P. 2012. Functional stem cell integration assessed by organotypic slice cultures. Curr Protoc Stem Cell Biol Chapter 2:Unit 2D 13. 10.1002/9780470151808.sc02d13s23.

Hofstetter AO, Saha S, Siljehav V, Jakobsson PJ and Herlenius E. 2007. The induced prostaglandin E2 pathway is a key regulator of the respiratory response to infection and hypoxia in neonates. Proc Natl Acad Sci U S A 104:9894-9899. 10.1073/pnas.0611468104.

Huckstepp RT, Bihi R, Eason R, Spyer KM, Dicke N, Willecke K, Marina N, Gourine AV and Dale N. 2010. Connexin hemichannel-mediated CO_2_-dependent release of ATP in the medulla oblongata contributes to central respiratory chemosensitivity. J Physiol 588:3901-3920. 10.1113/jphysiol.2010.192088.

Huckstepp RT, Eason R, Sachdev A and Dale N. 2010. CO2-dependent opening of connexin 26 and related beta connexins. J Physiol 588:3921-3931. 10.1113/jphysiol.2010.192096.

Huxtable AG, Zwicker JD, Alvares TS, Ruangkittisakul A, Fang X, Hahn LB, Posse de Chaves E, Baker GB, Ballanyi K and Funk GD. 2010. Glia contribute to the purinergic modulation of inspiratory rhythm-generating networks. J Neurosci 30:3947-3958. 30/11/3947 [pii] 10.1523/JNEUROSCI.6027-09.2010.

Lin W, Kemper A, McCarthy KD, Pytel P, Wang JP, Campbell IL, Utset MF and Popko B. 2004. Interferon-gamma induced medulloblastoma in the developing cerebellum. J Neurosci 24:10074-10083. 10.1523/JNEUROSCI.2604-04.2004.

Lorier AR, Huxtable AG, Robinson DM, Lipski J, Housley GD and Funk GD. 2007. P2Y1 Receptor Modulation of the Pre-Botzinger Complex Inspiratory Rhythm Generating Network In Vitro. Journal of Neuroscience 27:993-1005. 10.1523/jneurosci.3948-06.2007.

Meigh L, Greenhalgh SA, Rodgers TL, Cann MJ, Roper DI and Dale N. 2013. CO(2)directly modulates connexin 26 by formation of carbamate bridges between subunits. Elife 2:e01213. 10.7554/eLife.01213.

Okada Y, Sasaki T, Oku Y, Takahashi N, Seki M, Ujita S, Tanaka KF, Matsuki N and Ikegaya Y. 2012. Preinspiratory calcium rise in putative pre-Botzinger complex astrocytes. J Physiol 590:4933-4944. 10.1113/jphysiol.2012.231464.

Oku Y, Fresemann J, Miwakeichi F and Hulsmann S. 2016. Respiratory calcium fluctuations in low-frequency oscillating astrocytes in the pre-Botzinger complex. Respir Physiol Neurobiol 226:11-17. 10.1016/j.resp.2015.02.002.

Pascual O, Casper KB, Kubera C, Zhang J, Revilla-Sanchez R, Sul JY, Takano H, Moss SJ, McCarthy K and Haydon PG. 2005. Astrocytic purinergic signaling coordinates synaptic networks. Science 310:113-116. 10.1126/science.1116916.

Phillips WS, Herly M, Del Negro CA and Rekling JC. 2016. Organotypic slice cultures containing the preBotzinger complex generate respiratory-like rhythms. J Neurophysiol 115:1063-1070. 10.1152/jn.00904.2015.

Ruangkittisakul A, Kottick A, Picardo MC, Ballanyi K and Del Negro CA. 2014. Identification of the pre-Botzinger complex inspiratory center in calibrated “sandwich” slices from newborn mice with fluorescent Dbx1 interneurons. Physiol Rep 2. 10.14814/phy2.12111.

Ruangkittisakul A, Panaitescu B and Ballanyi K. 2011. K(+) and Ca(2)(+) dependence of inspiratory-related rhythm in novel “calibrated” mouse brainstem slices. Respir Physiol Neurobiol 175:37-48. 10.1016/j.resp.2010.09.004.

Ruangkittisakul A, Schwarzacher SW, Secchia L, Poon BY, Ma Y, Funk GD and Ballanyi K. 2006. High sensitivity to neuromodulator-activated signaling pathways at physiological [K+] of confocally imaged respiratory center neurons in on-line-calibrated newborn rat brainstem slices. J Neurosci 26:11870-11880. 26/46/11870 [pii] 10.1523/JNEUROSCI.3357-06.2006.

Schnell C, Fresemann J and Hulsmann S. 2011. Determinants of functional coupling between astrocytes and respiratory neurons in the pre-Botzinger complex. PLoS One 6:e26309. 10.1371/journal.pone.0026309.

Siljehav V, Hofstetter AM, Leifsdottir K and Herlenius E. 2015. Prostaglandin E2 Mediates Cardiorespiratory Disturbances during Infection in Neonates. J Pediatr 167:1207-1213 e1203. 10.1016/j.jpeds.2015.08.053.

Siljehav V, Olsson Hofstetter A, Jakobsson PJ and Herlenius E. 2012. mPGES-1 and prostaglandin E2: vital role in inflammation, hypoxic response, and survival. Pediatr Res 72:460-467. 10.1038/pr.2012.119.

Siljehav V, Shvarev Y and Herlenius E. 2014. Il-1beta and prostaglandin E2 attenuate the hypercapnic as well as the hypoxic respiratory response via prostaglandin E receptor type 3 in neonatal mice. J Appl Physiol (1985) 117:1027-1036. 10.1152/japplphysiol.00542.2014.

Smedler E, Malmersjo S and Uhlen P. 2014. Network analysis of time-lapse microscopy recordings. Front Neural Circuits 8:111. 10.3389/fncir.2014.00111.

Turovsky E, Karagiannis A, Abdala AP and Gourine AV. 2015. Impaired CO_2_ sensitivity of astrocytes in a mouse model of Rett syndrome. J Physiol 593:3159-3168. 10.1113/JP270369.

Turovsky E, Theparambil SM, Kasymov V, Deitmer JW, Del Arroyo AG, Ackland GL, Corneveaux JJ, Allen AN, Huentelman MJ, Kasparov S, Marina N and Gourine AV. 2016. Mechanisms of CO_2_/H+ Sensitivity of Astrocytes. J Neurosci 36:10750-10758. 10.1523/JNEUROSCI.1281-16.2016.

Young JK, Dreshaj IA, Wilson CG, Martin RJ, Zaidi SI and Haxhiu MA. 2005. An astrocyte toxin influences the pattern of breathing and the ventilatory response to hypercapnia in neonatal rats. Respir Physiol Neurobiol 147:19-30. 10.1016/j.resp.2005.01.009.

Young SZ, Platel JC, Nielsen JV, Jensen NA and Bordey A. 2010. GABA(A) Increases Calcium in Subventricular Zone Astrocyte-Like Cells Through L- and T-Type Voltage-Gated Calcium Channels. Front Cell Neurosci 4:8. 10.3389/fncel.2010.00008.

